# Inhibition of bladder cancer cell proliferation by 2-deoxy-D-glucose: A potential therapeutic approach

**DOI:** 10.1101/2025.11.09.687468

**Authors:** Xia Zhang, Rikiya Taoka, Akram Hossain, Yuki Matsuoka, Yoichiro Tohi, Hirohito Naito, Kazuya Akimitsu, Ken Izumori, Yoshiyuki Kakehi, Mikio Sugimoto

## Abstract

**Background:** Bladder cancer cells exhibit a high dependency on glycolysis for energy production and survival. 2-Deoxy-D-glucose (2-DG), a glucose analog, inhibits glycolysis by targeting hexokinase, leading to energy depletion and growth suppression in cancer cells.

**Objective:** This study investigated the effects of 2-DG on proliferation, glycolytic activity, oxidative stress, and apoptosis in human bladder cancer cells to evaluate its potential as a therapeutic agent.

**Methods:** Five human bladder cancer cell lines (RT112, 253J, T24, HT1197 and J82) were treated with increasing concentrations of 2-DG (0–20 mM) for up to 72 h. Cell viability was assessed by MTT assay, and glycolytic activity was evaluated by measuring hexokinase (HK) activity and intracellular ATP levels. Reactive oxygen species (ROS) production and apoptosis were analyzed by flow cytometry, and in vivo efficacy was examined using xenograft models in nude mice. All experiments were performed in triplicate.

**Results:** 2-DG significantly reduced cell viability in a dose-dependent manner across all cell lines, concomitant with decreased HK activity and ATP production (p < 0.05).

Treatment induced ROS accumulation and apoptosis, as evidenced by increased cleaved PARP and Bax expression and decreased p-Akt and Bcl-2 levels. In xenograft models, 2-DG treatment markedly suppressed tumor growth without apparent systemic toxicity.

**Conclusions:** 2-DG exerts potent antitumor effects against bladder cancer by inhibiting glycolysis, inducing oxidative stress, and activating apoptotic pathways. These findings support the therapeutic potential of 2-DG as a metabolic-targeting strategy for bladder cancer.

## Introduction

Bladder cancer is one of the most prevalent malignancies of the urinary tract and remains a major cause of cancer-related morbidity and mortality worldwide. Despite advances in surgery, chemotherapy, and immunotherapy, the prognosis of patients with advanced or recurrent bladder cancer remains poor. Therefore, new therapeutic strategies target fundamental metabolic pathways in cancer cells are urgently needed. Cancer cells are characterized by enhanced glycolysis even under normoxic conditions, a phenomenon known as the “Warburg effect.” This metabolic reprogramming enables rapid ATP generation and provides intermediates required for macromolecular biosynthesis, supporting uncontrolled cell growth and proliferation. Accordingly, inhibition of glycolysis has emerged as a promising approach for cancer therapy.

2-Deoxy-D-glucose (2-DG) is a glucose analog that competitively inhibits hexokinase, the first enzyme in the glycolytic pathway. After phosphorylation to 2-DG-6-phosphate, it cannot be further metabolized, resulting in glycolytic blockade, ATP depletion, and metabolic stress. Beyond glycolysis inhibition, 2-DG has also been shown to induce oxidative stress, disrupt N-linked glycosylation, and modulate autophagy and apoptosis in various cancer types. These multifaceted actions make 2-DG an attractive candidate for targeting cancer metabolism.

Previous studies demonstrated that 2-DG suppresses proliferation and enhances chemosensitivity in several cancers, including glioma, breast, and prostate cancers. It’s inhibitory effects on bladder cancer cells have also been reported (PMID: 36765947; PMID: 26504012). However, the precise mechanisms underlying 2-DG-mediated cytotoxicity in bladder cancer, particularly in relation to glycolysis inhibition, oxidative stress, and apoptotic signaling, remain incompletely understood.

The present study aimed to investigate the antitumor effects and underlying mechanisms of 2-DG in bladder cancer. We examined its impact on cell viability, glycolytic activity, ROS generation, and apoptotic signaling in multiple bladder cancer cell lines, as well as its therapeutic efficacy in a xenograft model. Our findings provide new insights supporting 2-DG as a potential metabolism-targeted therapeutic agent for bladder cancer.

## Materials and Methods

### Cell Lines

Five human bladder cancer cell lines (253 J, HT1197, J82, RT112, and T24) were obtained from the Japanese distributor of ATCC®. Cells were cultured in RPMI 1640 medium supplemented with 10% fetal bovine serum and 1% penicillin-streptomycin and maintained at 37°C in a humidified atmosphere containing 5% CO₂.

### Cell Viability Assay

Cell viability was measured using the 3-(4,5-dimethylthiazol-2-yl)-2,5-diphenyltetrazolium bromide (MTT) assay. Cells were seeded at 4,000 cells/well in 96-well plates and allowed to attach for 24 h. Cells were treated with 10, 25, or 50 mM 2-DG for 24 h. Cell viability was determined using the Cell Proliferation Kit I (Roche, Mannheim, Germany). Optical density was measured at 570 nm with a reference wavelength of 750 nm. All treatments were performed in triplicate.

### Hexokinase (HK) Activity Assay

Hexokinase activity was determined using the Hexokinase Activity Assay Kit (Cell Biolabs, Inc.). RT112 and 253 J cells were seeded in 6-well plates at 4,000 cells/mL. After 24 h, cells were treated with 10, 25, or 50 mM 2-DG for 24 h. Supernatants and cell lysates were prepared according to the manufacturer’s instructions. Hexokinase activity was measured at 450 nm using a microplate reader. All assays were performed in triplicate.

### Cellular ATP Level Assay

Cellular ATP levels were measured using the ATP Colorimetric Assay Kit (BioVision, Milpitas, CA, USA). RT112 and 253 J cells were seeded at 4,000 cells/mL in 6-well plates and treated with 2-DG for 24 h. After treatment, cells were lysed, and ATP standards and samples were prepared in a 96-well plate. Reactions were initiated by adding the reaction mixture and incubating for 30 min at room temperature in the dark.

Absorbance was measured at 570 nm using a microplate reader. All assays were performed in triplicate.

### Intracellular ROS Assay

Intracellular reactive oxygen species (ROS) levels were evaluated using DCF-DA and flow cytometry. Cells were plated at 3 × 10⁵ cells/cm² in 5 mm culture dishes. After 24 h, cells were treated with methanol (control) or various concentrations of 2-DG for 1 h. Cells were then incubated with 10 μM DCF-DA for 30 min at 37°C. ROS levels were measured using a CytoFLEX S flow cytometer (Beckman Coulter, CA, USA) and analyzed with CytExpert software.

### Western Blotting Analysis

After 48 h of 2-DG treatment, cells were harvested and lysed using Cell Lysis Buffer M (Wako Pure Chemical Industries, Ltd., Osaka, Japan) according to the manufacturer’s instructions. Primary antibodies included PARP, cleaved PARP, Akt, Bax, and Bcl-2 (Abcam, Tokyo, Japan; 1:1000) and GAPDH (AC74, Sigma-Aldrich, Tokyo, Japan; 1:2000) as a loading control. Membranes were incubated with HRP-conjugated secondary antibodies for 1 h, and signals were detected using a chemiluminescence kit (Amersham Biosciences).

### Xenograft Tumor Model in Nude Mice

Male BALB/cA Jcl-nu/nu mice (6 weeks old) were purchased from Charles River Laboratories Japan, Inc. (Yokohama, Japan) and maintained in accordance with the Institutional Regulations for Animal Experiments at Kagawa University (ethical permit no. 17631). RT112 and 253 J cells (1 × 10⁷) were prepared in collagen gel and injected subcutaneously into the backs of the mice. When tumor volume reached approximately 100 mm³, mice were randomly divided into three groups (six mice per group) and treated intraperitoneally with PBS (control) or 2-DG (400 mg/kg) 5 days per week for 32 days. Tumor size was measured every 4 days using calipers and calculated as: Tumor volume=length×(width)2×0.5¥text{Tumor volume} = ¥text{length} ¥times (¥text{width})^2 ¥times 0.5Tumor volume=length×(width)2×0.5 At the endpoint, tumors, kidneys, and liver were collected, fixed in paraffin, sectioned (4 μm), and stained to evaluate tumor proliferation.

### Statistical Analysis

Data are presented as mean ± standard deviation (SD). Comparisons between two groups were performed using an unpaired Student’s t-test. Statistical significance was defined as p < 0.05. All experiments were performed in at least three independent replicates.

## Results

### 2-DG Inhibits Bladder Cancer Cell Viability and Glycolytic Activity

To evaluate the cytotoxic effects of 2-DG on bladder cancer cells, five human bladder cancer cell lines (RT112, 253 J, T24, HT1197, and J82) were treated with increasing concentrations of 2-DG for 24 h, and cell viability was assessed by the MTT assay. As shown in **Fig 1A**, 2-DG significantly reduced cell viability in a dose-dependent manner across all cell lines. At 50 mM, 2-DG decreased viability by 21.4 ± 1.2% in RT112, 35.2 ± 2.2% in 253 J, 13.1 ± 1.2% in T24, 38.5 ± 2.1% in HT1197, and 47.9 ± 3.4% in J82 cells relative to untreated controls (p < 0.05).

**Fig 1.**
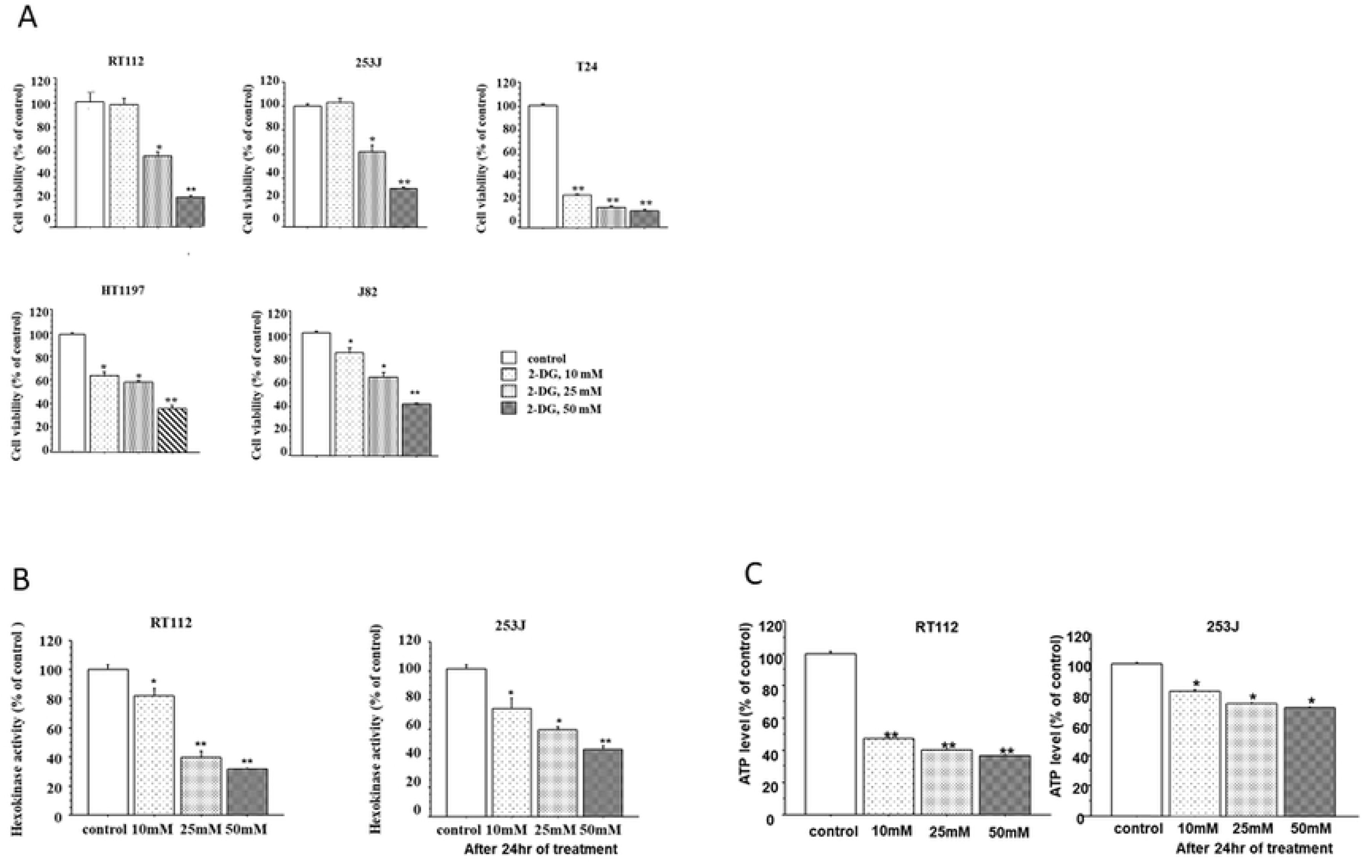
2-DG inhibits bladder cancer cell viability and glycolysis. (A) Cell viability of five human bladder cancer cell lines (RT112, 253 J, T24, HT1197, and J82) after 24 h of 2-DG treatment measured by MTT assay. (B) Hexokinase (HK) activity in RT112 and 253 J cells after 24 h of 2-DG treatment. (C) Cellular ATP levels in RT112 and 253 J cells after 24 h of 2-DG treatment. Data are mean ± SD from three independent experiments. *p < 0.05, **p < 0.005.

To determine whether this cytotoxic effect was associated with glycolytic inhibition, hexokinase (HK) activity was measured after 24 h of treatment. As shown in **Fig 1B**, 2-DG induced a marked, dose-dependent reduction in HK activity in both RT112 and 253 J cells, with decreases of 37% and 48%, respectively, at 50 mM (p < 0.005). Similarly, intracellular ATP levels were significantly reduced in a concentration-dependent manner (**Fig 1C**), with 50 mM 2-DG reducing ATP by 38% in RT112 and 48% in 253 J cells. Collectively, these findings indicate that 2-DG effectively suppresses bladder cancer cell proliferation by disrupting glycolytic enzyme activity and cellular energy metabolism.

### 2-DG Induces Oxidative Stress in Bladder Cancer Cells

Because inhibition of glycolysis can cause redox imbalance, we next assessed intracellular reactive oxygen species (ROS) production using DCF-DA staining followed by flow cytometric analysis. Treatment of RT112 and 253 J cells with 2-DG for 1 h led to a significant and dose-dependent increase in ROS generation compared with controls (**Fig 2**). Exposure to 50 mM 2-DG elevated ROS levels by 172.1 ± 5.6% in RT112 and 141 ± 2.5% in 253 J cells (p < 0.005). These results demonstrate that 2-DG induces oxidative stress, suggesting that ROS accumulation contributes to its cytotoxic effects in bladder cancer cells.

**Figure 2.**
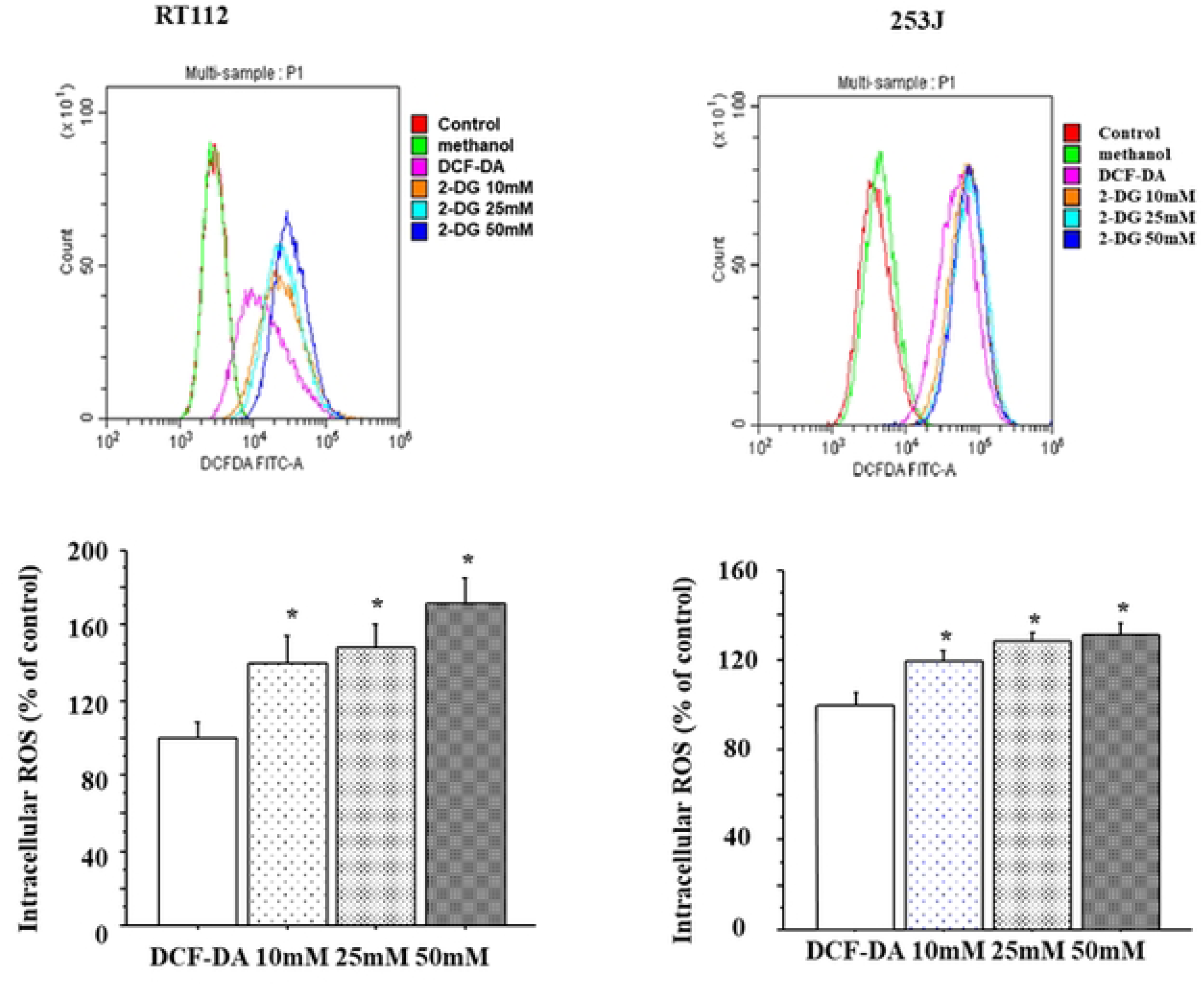
2-DG induces oxidative stress in bladder cancer cells. Intracellular reactive oxygen species (ROS) levels in RT112 and 253 J cells after 1 h of 2-DG treatment, measured using DCF-DA and flow cytometry. Data are mean ± SD of three independent experiments. Statistically significant increases compared with control are indicated by *p < 0.05 and **p < 0.005.

### 2-DG Activates Apoptotic Signaling Pathways

To elucidate whether 2-DG-induced cell death involves apoptosis, the expression of apoptosis-related proteins was examined by western blotting. Following 48 h of 2-DG treatment, a concentration-dependent downregulation of full-length PARP and phosphorylated Akt (p-Akt) was observed, accompanied by an increase in cleaved PARP levels in both RT112 and 253 J cells (**Fig 3A**). These changes are indicative of caspase-dependent apoptotic signaling activation.

**Figure 3.**
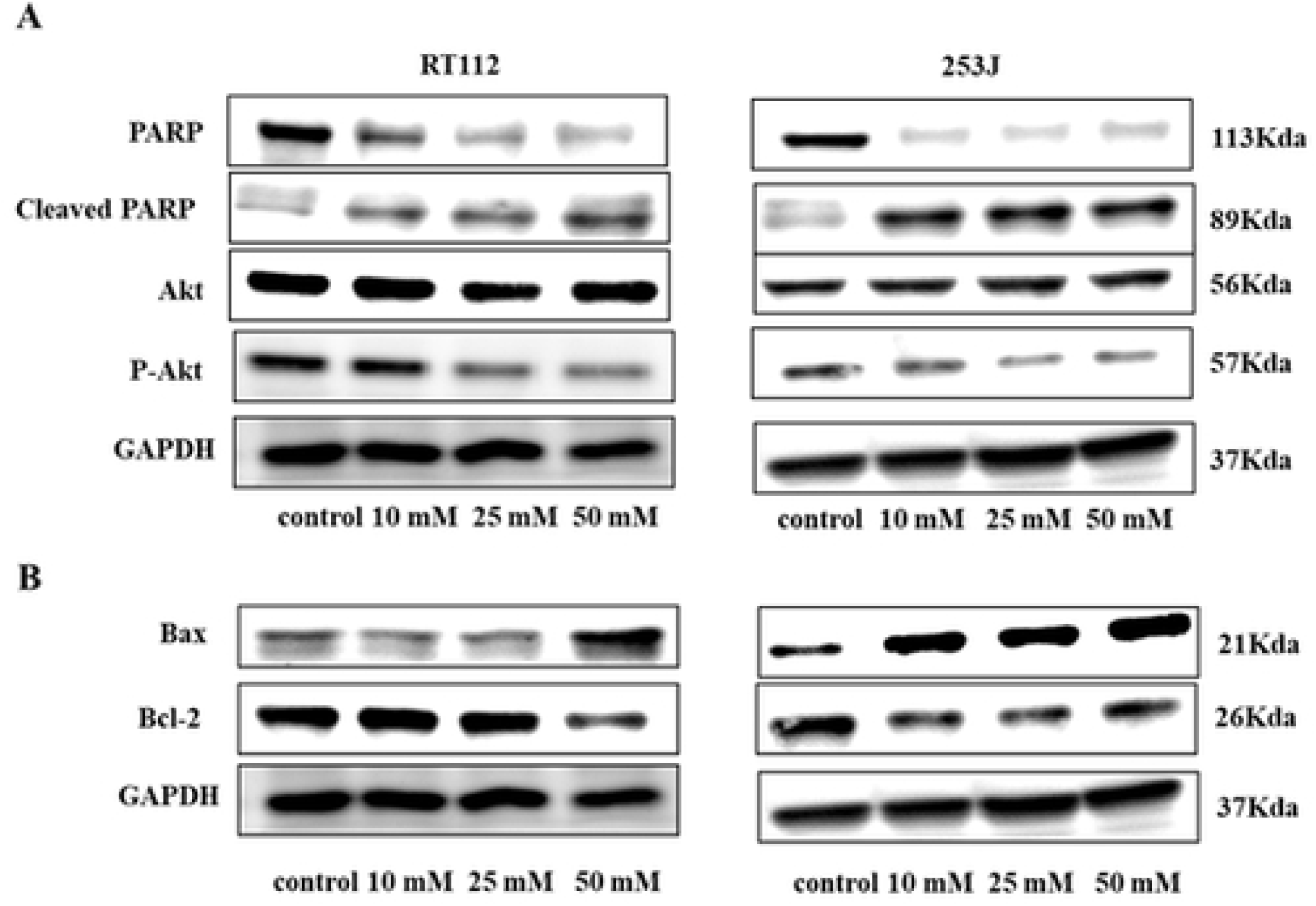
2-DG modulates apoptotic pathways in bladder cancer cells. Expression of PARP, cleaved PARP, and p-Akt proteins in RT112 and 253 J cells 48 h post-2-DG treatment. (B) Expression of Bax and Bcl-2 proteins under the same conditions. GAPDH was used as a loading control. Data from three independent experiments show dose-dependent induction of apoptosis via mitochondrial pathways.

Furthermore, 2-DG altered mitochondrial apoptotic regulators (**Fig 3B**). The expression of the pro-apoptotic protein Bax increased, whereas the anti-apoptotic protein Bcl-2 decreased with increasing 2-DG concentrations, resulting in an elevated Bax/Bcl-2 ratio. These findings suggest that 2-DG induces apoptosis through both inhibition of the Akt survival pathway and activation of the mitochondrial apoptotic cascade.

### 2-DG Suppresses Tumor Growth In Vivo Without Systemic Toxicity

The antitumor efficacy of 2-DG was next evaluated in vivo using RT112 and 253 J xenograft models. Mice were administered 2-DG intraperitoneally for 32 days, and tumor growth was monitored throughout the treatment period. As shown in **Fig 4A**, tumor volumes were significantly reduced in 2-DG–treated groups compared with PBS controls (p < 0.005).

**Figure 4.**
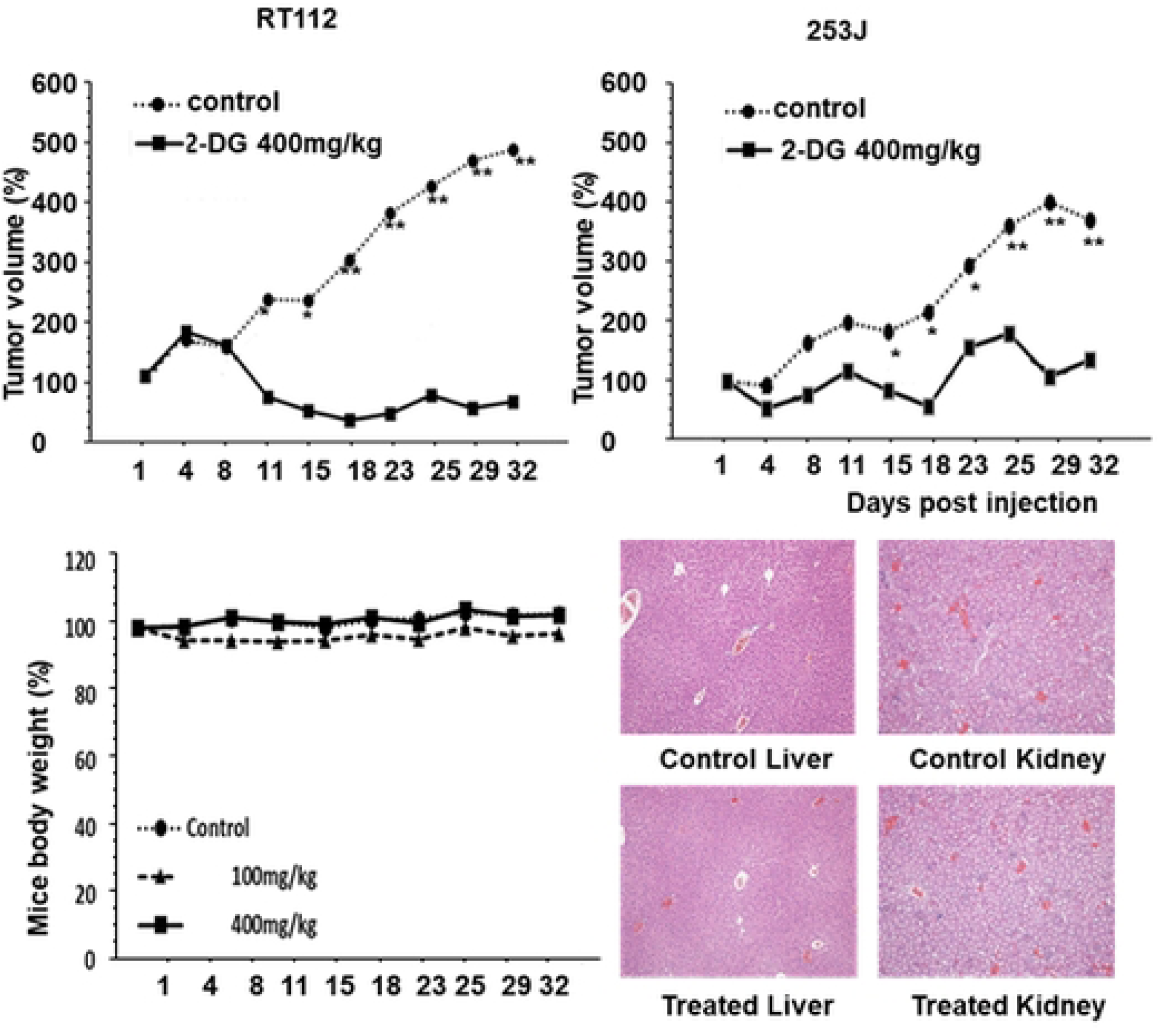
2-DG suppresses tumor growth in bladder cancer xenografts. (A) Tumor volumes of RT112 and 253 J xenografts in nude mice after 32 days of intraperitoneal 2-DG treatment or PBS control. (B) Body weight changes and histological evaluation of kidneys and livers from treated mice. Data are mean ± SD. Statistically significant differences compared with control are indicated by *p < 0.05 and **p < 0.005. No significant systemic toxicity was observed at the tested dose.

Importantly, 2-DG treatment did not cause any noticeable systemic toxicity. No significant changes in body weight were observed (**Fig 4B**), and no behavioral abnormalities were noted during the study period. Histological examination of kidney and liver tissues revealed no pathological alterations (**Fig 4C**), indicating that 2-DG exerts its antitumor effects without inducing major organ toxicity.

Collectively, these results demonstrate that 2-DG effectively inhibits bladder cancer growth both in vitro and in vivo by impairing glycolysis, inducing oxidative stress, and activating apoptotic signaling pathways.

## Discussion

In this study, we demonstrated that 2-deoxy-D-glucose (2-DG) effectively inhibits the viability of bladder cancer cells in a dose-dependent manner, consistent with previous reports describing its antiproliferative effects in various cancer types [2,7,18]. One of the key mechanisms underlying the anti-tumor effects of 2-DG is the inhibition of glycolysis. Our results showed that 2-DG treatment caused a dose-dependent reduction in hexokinase activity, the rate-limiting enzyme in glycolysis, which plays a critical role in cancer cell metabolism. This finding aligns with previous studies demonstrating 2-DG-mediated HK inhibition in different cancer models [8,9]. By targeting HK, 2-DG disrupts the glycolytic pathway, leading to decreased ATP production, which is essential for cancer cell survival and proliferation. Indeed, our results also revealed a significant reduction in intracellular ATP levels following 2-DG treatment, further supporting its glycolytic inhibition. Similar reductions in ATP have been reported in various cancer cell lines treated with 2-DG [10,11].

In addition to its metabolic effects, 2-DG induces oxidative stress in bladder cancer cells, as evidenced by a significant increase in intracellular ROS levels. Elevated ROS can cause cytotoxic damage and trigger apoptosis, suggesting that oxidative stress contributes to the antitumor activity of 2-DG [12,13].

Furthermore, 2-DG modulated key apoptotic markers. Treatment resulted in downregulation of total PARP expression and upregulation of cleaved PARP, accompanied by decreased phosphorylated Akt (p-Akt) levels, indicating activation of apoptotic signaling. We also observed a dose-dependent increase in Bax and a corresponding decrease in Bcl-2 protein levels, highlighting the involvement of the mitochondrial apoptotic pathway. It should be noted that our study evaluated steady-state protein expression rather than transcriptional changes. These findings are consistent with earlier reports demonstrating that 2-DG regulates apoptosis in various cancer models [14,15].

In vivo, 2-DG significantly suppressed tumor growth in bladder cancer xenografts without affecting body weight or histological features of the kidney and liver, suggesting minimal systemic toxicity at the tested dose [16]. However, this study has certain limitations. In vitro assays and nude mouse xenograft models have limited predictive value for clinical outcomes, and further preclinical and clinical investigations are required to confirm the therapeutic efficacy of 2-DG in bladder cancer.

In summary, our findings indicate that 2-DG exerts potent antitumor effects by inhibiting glycolysis, inducing oxidative stress, and modulating apoptotic signaling pathways. These results provide a mechanistic evidence supporting the potential development of 2-DG as a metabolic-targeting therapeutic strategy for bladder cancer.

## Supplementary Materials

Not applicable.

## Author Contributions

Conceptualization: Xia Zhang, Rikia Taoka, Yoshiyuki Kakehi.

Data curation: Xia Zhang, Yoichiro Tohi.

Formal analysis: Rikia Taoka, Xia Zhang.

Funding acquisition: Rikia Taoka.

Investigation: Xia Zhang, Rikia Taoka, Yoichiro Tohi, Yuki Matsuoka, Yoshiyuki Kakehi.

Supervision: Mikio Sugimoto.

Visualization: Xia Zhang.

Writing-original draft: Xia Zhang.

Writing-review and editing: Rikia Taoka, Akram Hossain, Xia Zhang, Yuki Matsuoka, Yoichiro Tohi, Mikio Sugimoto, Yoshiyuki Kakehi.

All authors have read and agreed to the published version of the manuscript.

## Funding

Not applicable

## Institutional Review Board Statement

Not applicable

## Informed Consent Statement

Not applicable

## Data Availability Statement

Not applicable

## Acknowledgments

Not applicable

## Conflicts of Interest

The authors declare no conflict of interest.

## Notes

### Competing Interest Statement

The authors have declared no competing interest.

